# Insecticidal and Feeding Deterrent Effects of Menadione and Menadione Sodium Bisulfite in Wood Treatment against the Formosan Subterranean Termite

**DOI:** 10.1101/2025.09.19.677127

**Authors:** Arjun Khadka, Myriel Green, Roger A. Laine, Qian Sun

## Abstract

The Formosan subterranean termite, *Coptotermes formosanus* Shiraki, is an invasive species and among the most economically damaging structural pest worldwide. Various insecticides and natural substances have been investigated for their management. Among these, naphthoquinones have demonstrated insecticidal activity against termites and diverse other insect pests. Menadione (vitamin K_3_), a naphthoquinone derivative, has been examined for its toxicity and repellency against *C. formosanus* when applied to soil, the tunneling substrate for termites. However, its potential for wood preservation was not investigated. In this study, we tested the water-insoluble menadione and its water-soluble derivative, menadione sodium bisulfite (MSB), for their effects on mortality and food consumption in *C. formosanus* through wood treatment over 28 days. Our results revealed toxicity for both compounds, with significant termite mortality observed between 500 and 1000 ppm in a no-choice assay. In a choice assay where termites were provided with treated and untreated wood, menadione at concentrations ≥ 500 ppm exhibited a strong feeding deterrent effect and reduced wood damage. However, such effects were not consistently detected for MSB ranging from 1 to 1000 ppm. These findings suggest that menadione is more effective than MSB and has potential as a wood preservation agent.

## 1. Introduction

Subterranean termites (Blattodea: Heterotermitidae) account for approximately 80% of the global economic impact of all termite pests, causing an estimated USD 32 billion per year in treatment and structural repair cost [1]. Among them, the invasive Formosan subterranean termite, *Coptotermes formosanus* Shiraki, is one of the most economically damaging termites in the world [1]. Native to eastern Asia [2], this termite is a serious structural pest in Hawaii and the southeastern United States, with annual losses of USD 1 billion annually estimated in 2005 [3]. This cost has likely increased substantially in recent years due to rising management and structural repair expenses, as well as the continued spread of the termite [4-7].

A variety of termiticides have been investigated and applied to termite management practices, including soil treatment, baiting, and wood preservation [8-10]. Liquid termiticides are among the conventional and most widely used termite control methods, and they are typically applied to the soil around and under the protected structure. These termiticides are categorized into repellents and non-repellents. Compared to repellents, the use of non-repellent termiticides, including fipronil, imidacloprid, chlorfenapyr, and chlorantraniliprole, has been validated to be more effective, causing higher termite mortality and more successful protection [8, 11]. Repellent termiticides, mainly pyrethroids, pose challenges because termites avoid treated areas and thorough treatment of all entry points is required; therefore, these compounds are better suited for preventive applications [8, 11, 12]. While repellency may limit their use in soil barrier treatments, chemicals with repellent or feeding deterrent effects can be used as wood preservatives instead. For example, bifenthrin and permethrin, which are repellent pyrethroids, are effectively used in the wood preservation industry [13-17]. Additionally, many naturally derived compounds with insecticidal, repellent, or feeding deterrent activities have been explored, offering alternative options for termite control [18-20].

Menadione (2-methyl-1, 4-naphthoquinone), also known as vitamin K_3_, is a water-insoluble compound and a precursor to biologically active forms of vitamin K [21]. This compound has a low level of mammalian toxicity (0.5 g/kg orally in mice) [22]. Although not allowed in human food, both menadione and its water-soluble derivative, sodium menadione bisulfite (MSB), have been used as an inexpensive nutritional supplement in animal feeds [23]. These synthetic naphthoquinone derivatives and naturally occurring naphthoquinones show insecticidal properties against various urban and agricultural pests, suggesting their use as safe and cost-efficient pest control agents [24]. For example, plumbagin (5-hydroxy-2-methyl-1,4-naphthoquinone) has demonstrated strong acute toxicity against the house fly (*Musca domestica* Linnaeus) [25]. 5-Hydroxy-2-methyl-1,4-naphthoquinone isolated from the roots of persimmon (*Diospyros kaki* Thunberg), as well as its structural derivatives, were reported to have potent larvicidal activities against three mosquitoes (*Aedes aegypti* L., *Culex pipiens* L., and *Ochlerotatus togoi* Theobald) [26]. Similarly, MBS exhibited insecticidal properties against the sweet potato whitefly *Bemisia tabaci* (Gennadius) and the tomato psyllid *Bactericera cockerelli* (Sulc) [27]. In addition, both toxic and repellent effects of naphthoquinones extracted from the heartwood of teak (*Tectona grandis* L.f.) have been demonstrated on the drywood termite *Incisitermes marginipennis* (Latreille) [28]. Menadione and other naphthoquinones presumably have a mode of action to disrupt mitochondrial respiration in insects, as evidenced by their mitochondrial oxidative phosphorylation uncoupling and cytotoxic activities in mammalian cells [29, 30].

Our previous study evaluated the effects of menadione on *C. formosanus* through soil treatment, and the results showed both strong toxic and repellent activities [31]. When menadione at 6-600 ppm was applied to sand, 90-100% mortality in seven days was observed in a no-choice assay; in a choice assay where termites were exposed to treated and untreated sand, menadione at ≥ 6 ppm showed significant repellent activity and prevented termites from moving to the treated side for feeding [31]. Although repellency is a less desirable property for soil termiticides than non-repellency, investigating menadione for its potential use as a wood preservative remains valuable. In this study, we examined both menadione and MSB for their influences on the mortality and feeding of *C. formosanus* termites over a 28-day period through wood treatment. First, a no-choice assay was conducted to evaluate the toxicity and the effects on food consumption of wood treated with each compound at concentrations ranging from 0 to 1000 ppm. Additionally, a choice assay comparing untreated and treated wood was conducted to determine the potential feeding deterrent effects of menadione and MSB.

## 2. Materials and Methods

### 2.1 Insects

Workers and soldiers from *C. formosanus* colonies were collected from Brechtel Park, New Orleans, Louisiana, using pine wood traps buried in the ground [32]. The collected termites were then maintained in laboratory at 25 ± 1 °C and kept in complete darkness in acrylic plastic containers (45.7 × 30.4 × 22.8 cm^3^, Rubbermaid, Atlanta, GA). Organic soil (Miracle-Gro All Purpose for In-Ground Use, Scotts Miracle-Gro, Marysville, OH, USA) was placed at the bottom, and moistened pine wood blocks were provided as the food source. The termites were used for experimentation after a two-week acclimation period in the laboratory.

### 2.2 No-Choice Assay

A no-choice assay was conducted to test if wood treated with either menadione or MSB at different concentrations affected the mortality and food consumption of *C. formosanus* termites. Kiln-dried pine wood blocks (*Pinus* sp., purchased from the local Home Depot store) were cut into pieces (3.6 × 1.7 × 0.6 cm^3^), oven-dried at 100 °C for 24 hours, and weighed. The wood pieces were then immersed in solutions of menadione or MSB (≥ 98.0% and ≥ 95.0%, respectively, MilliporeSigma, Burlington, MA, USA) at concentrations of 0 (solvent control), 1, 10, 100, 500, and 1000 ppm for 30 seconds. The concentrations were selected based on a previous study [31]. Menadione solutions were prepared with acetone as the solvent, while MSB solutions were prepared with distilled water. For the menadione treatment, the immersed wood pieces were allowed to dry in the fume hood for 24 hours to facilitate acetone evaporation before use for experiments.

The assay was conducted in round plastic containers (3.6 cm in height and 5.0 cm in diameter, Pioneer Plastics, North Dixon, KY, USA) (Figure 1A). Each container was filled with 60 g of sand (Quikrete Premium Play Sand, Atlanta, GA, USA), and 9 mL of distilled water was added to reach 15% moisture. To prevent any leaching into the sand, each treated wood piece was placed on an aluminum foil sheet (4.1 cm in length and 2.5 cm in width), which was then positioned on top of the sand. A group of 100 termites (90 workers and 10 soldiers) was released into each container, which was then covered with a lid. The experimental containers were placed inside a larger sealed plastic container (25.4 × 17.8 × 7.6 cm^3^), where moistened paper towels were provided at the bottom to maintain about 98% relative humidity (RH). The experiment was conducted for 28 days at room temperature (25 ± 1 °C) in complete darkness. At the end of the assay, termite mortality was recorded, and food consumption was calculated based on the weight difference of the wood after oven-drying at 100 °C for 24 hours. Termites from three *C. formosanus* colonies were used, with each colony replicated three times using different groups of termites.

**Figure 1.**
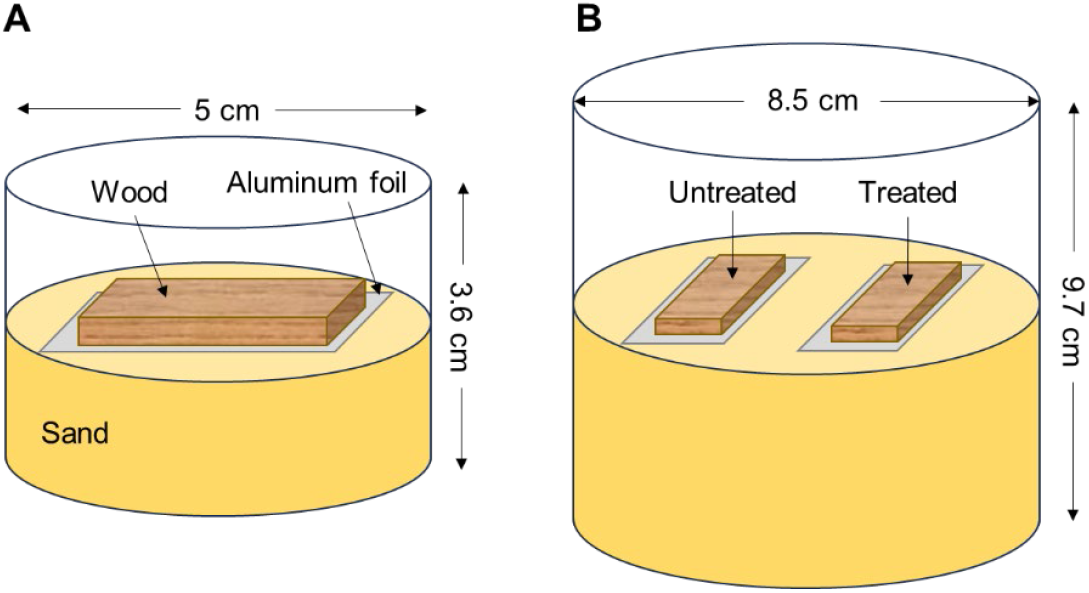
Schematic illustration of experimental setups. (**A**) No-choice assay; (**B**) Choice assay.

### 2.3 Choice Assay

A choice assay was conducted to determine if menadione or MSB exhibited any deterrent effects on *C. formosanus* termites through wood treatment. The assays were conducted in glass jars (9.7 cm in height and 8.5 cm in diameter, Qorpak, Clinton, PA, USA) (Figure 1B). Each jar contained 350 g of sand with 52.5 mL of distilled water added to reach 15% moisture. Menadione and MSB solutions at concentrations of 0 (solvent control), 1, 10, 100, 500, and 1000 ppm were prepared to treat pine wood pieces as described above. In each jar, two wood pieces (one untreated and one treated) were placed on top of the sand, each held separately by an aluminum foil sheet (4.1 cm in length and 2.5 cm in width) to prevent leaching into the sand. A group of 100 termites (90 workers and 10 soldiers) was then introduced to each jar. The jars were covered with a lid, slightly loosened to allow ventilation.

The experimental glass jars were placed in a larger sealed container (31.8 × 25.6 × 9.8 cm^3^) with moistened paper towel at the bottom to maintain about 98% RH. The experiment was conducted for 28 days at room temperature (25 ± 1 °C) in complete darkness. At the end of the assay, termite mortality was recorded, and food consumption of each wood piece was calculated by pre-and post-experiment weight difference of the wood, which was oven-dried at 100 °C for 24 hours. Termites from three *C. formosanus* colonies were used, with each colony replicated three times using different groups of termites.

### 2.4 Data Analysis

Data were analyzed using the R software v4.3.2 (the R Foundation, Vienna, Austria) [33] and visualized using PRISM v10.1.2 (GraphPad Software, San Diego, CA, USA). Since assays for menadione and MSB were conducted at different times with slightly different treatment methods, the data for the two compounds were analyzed separately. Additionally, in the no-choice assays, data from several replicates were excluded due to fungal growth in the early stages, resulting in varying sample sizes across treatments (menadione: *n* = 8 for 0, 500 and 1000 ppm, *n* = 7 for 1 ppm, and *n* = 9 for 10 and 100 ppm; MSB: *n* = 8 for 0 ppm, *n* = 7 for 1 ppm, and *n* = 9 for all other treatments). For all data, normality was tested using Shapiro-Wilk tests, and homogeneity of variance was evaluated using Levene’s test (α = 0.05). As data did not meet the assumptions for parametric analyses, a generalized linear mixed model (GLMM) was used to analyze the mortality data from all assays and the food consumption data from the no-choice assays. Colony was coded as a random factor and treatment as a fixed factor, and GLMM was followed by mean separation tests using the *lsmeans* package. Wilcoxon signed-rank test was performed for pairwise comparisons of food consumption data in the choice assays. The original data are provided in the supplemental materials (Table S1).

## 3. Results

### 3.1. No-Choice Assay

Menadione affected the survival and food consumption of *C. formosanus* termites through wood treatment. At 500 ppm, significantly higher termite mortality (35.88 ± 11.18%, mean ± SE) compared to the control (0 ppm, 9.62 ± 1.94%) was observed (GLMM, *df* = 41, *p* < 0.05, Figure 2A). Although mortality at 1000 ppm (27.50 ± 10.41%) was higher than that at 0, 1, 10, and 100 ppm, the differences among these groups were not statistically significant (GLMM, *df* = 41, *p* > 0.05, Figure 2A). Additionally, termites consumed significantly less wood treated with 1000 ppm of menadione (0.06 ± 0.02 g) compared to the control (0.23 ± 0.05 g) and concentrations ranging from 1 to 500 ppm (GLMM, *df* = 41, *p* < 0.05, Figure 2B).

**Figure 2.**
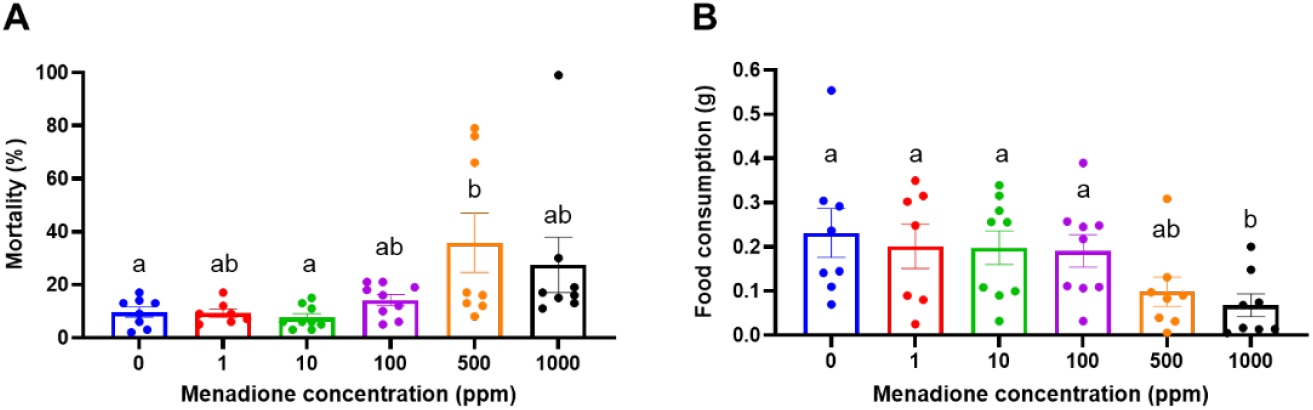
Effects of menadione on termite mortality and feeding in the no-choice assay. (**A**) Termite mortality after 28 days of treatments. (**B**) Wood consumption after 28 days of treatments. Groups sharing the same letter are not significantly different (GLMM, *df* = 41, *p* > 0.05). Bars represent mean ± SE, and dots show individual values.

Similar to menadione, MSB affected termite mortality according to concentration. At 1000 ppm, significantly higher termite mortality (75.56 ± 10.42%) was observed compared to the 0 ppm control (12.13 ± 3.46%) (GLMM, *df* = 43, *p* < 0.05, Figure 3A). Termite mortality did not significantly differ between the control and MSB treatment at concentrations from 1 to 500 ppm (GLMM, *df* = 43, *p* > 0.05, Figure 3A). Food consumption at 1000 ppm of MSB (0.09 ± 0.04 g) was lower than that in the control group (0.19 ± 0.04 g); however, no significant differences were observed between these groups or across other tested concentrations (GLMM, *df* = 43, *p* > 0.05, Figure 3B). Representative images of wood damage observed in the no-choice assays for menadione and MSB are presented in Figure 4.

**Figure 3.**
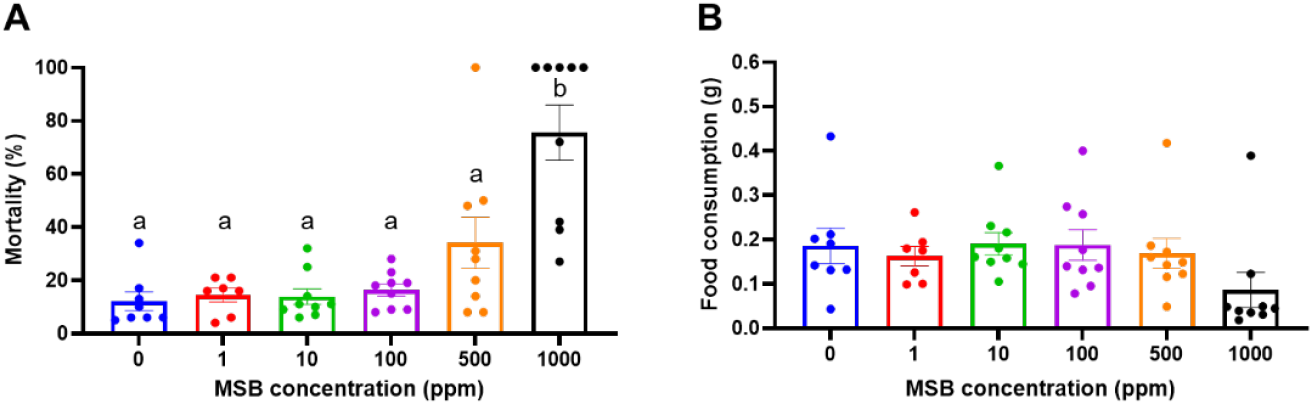
Effects of MSB on termite mortality and feeding in the no-choice assay. (**A**) Termite mortality 28 days post-treatment. Groups sharing the same letter are not significantly different (GLMM, *df* = 43, *p* > 0.05). (**B**) Wood consumption over 28 days post-treatment. No significant differences were detected among groups (GLMM, *df* = 43, *p* > 0.05). Bars represent mean ± SE, and dots show individual values.

**Figure 4.**
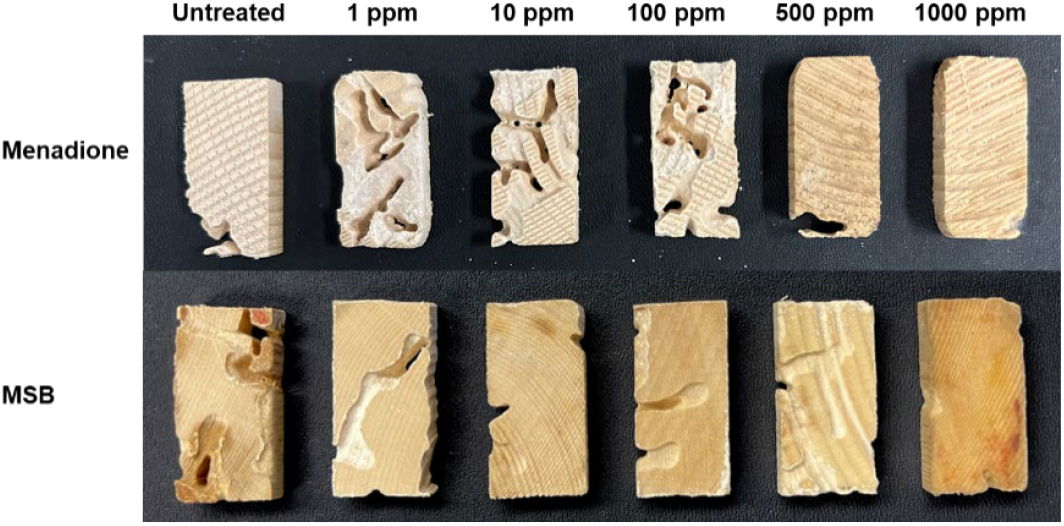
Representative images of wood damage by termites 28 days post-menadione and MSB treatments in no-choice assays.

### 3.2 Choice Assay

When termites were provided with both an untreated and a menadione-treated wood piece, overall mortality did not differ significantly across the tested concentrations (GLMM, *df* = 46, *p* > 0.05, Figure 5A), with mean mortality below 18% in each group. Food consumption did not differ significantly between untreated and treated wood at menadione concentrations ≤ 100 ppm (Wilcoxon signed-rank test, *p* > 0.05, Figure 5B). However, consumption was significantly reduced for wood treated with menadione at 500 and 1000 ppm (0.04 ± 0.01 and 0.03 ± 0.01 g, respectively) compared to the paired untreated pieces (0.21 ± 0.05 and 0.18 ± 0.04 g, respectively) (Wilcoxon signed-rank test, *p* < 0.01, Figure 5B).

**Figure 5.**
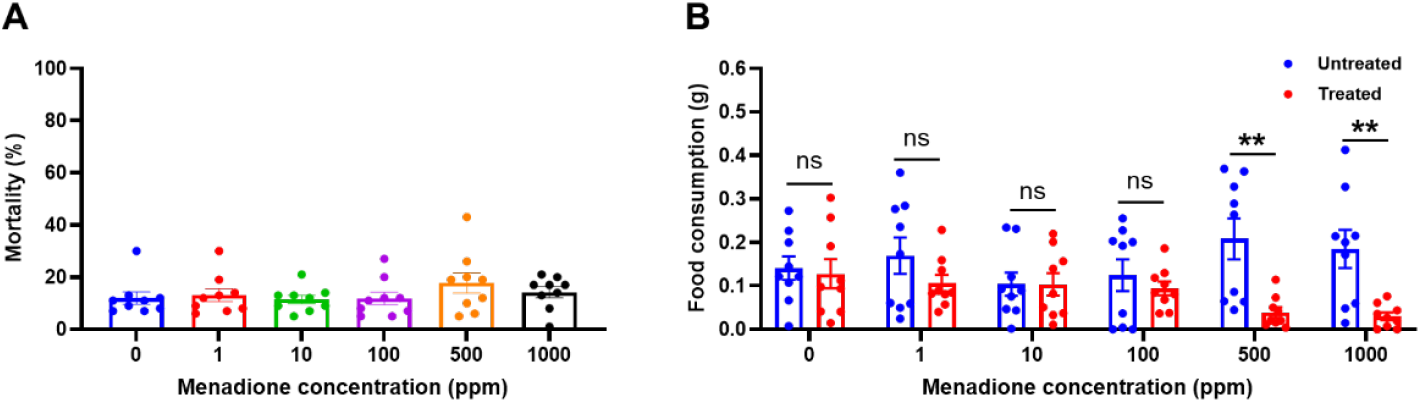
Effects of menadione on termite mortality and feeding in the choice assay. (**A**) Termite mortality 28 days post-treatment. No significant differences were detected among groups (GLMM, *df* = 46, *p* > 0.05). (**B**) Wood consumption over 28 days post-treatment (Wilcoxon signed-rank test; ns: not significant (*p* > 0.05); ** *p* < 0.01). Bars represent mean ± SE, and dots show individual values.

In the choice assay with MSB, overall termite mortality was significantly higher at the concentration of 1000 ppm (48 ± 10.39%) than other groups (GLMM, *df* = 46, *p* < 0.05), while no significant differences were found among the other concentrations from 0 to 500 ppm (GLMM, *df* = 46, *p* > 0.05, Figure 6A), where mean mortality remained below 15%. A significant reduction in the consumption of treated wood compared to untreated wood was observed only at 100 ppm (Wilcoxon signed-rank test, *p* < 0.05), while no significant differences in consumption of paired wood pieces were found at other concentrations (Wilcoxon signed-rank test, *p* > 0.05, Figure 6B). Representative images of wood damage observed in the choice assays for menadione and MSB are presented in Figure 7.

**Figure 6.**
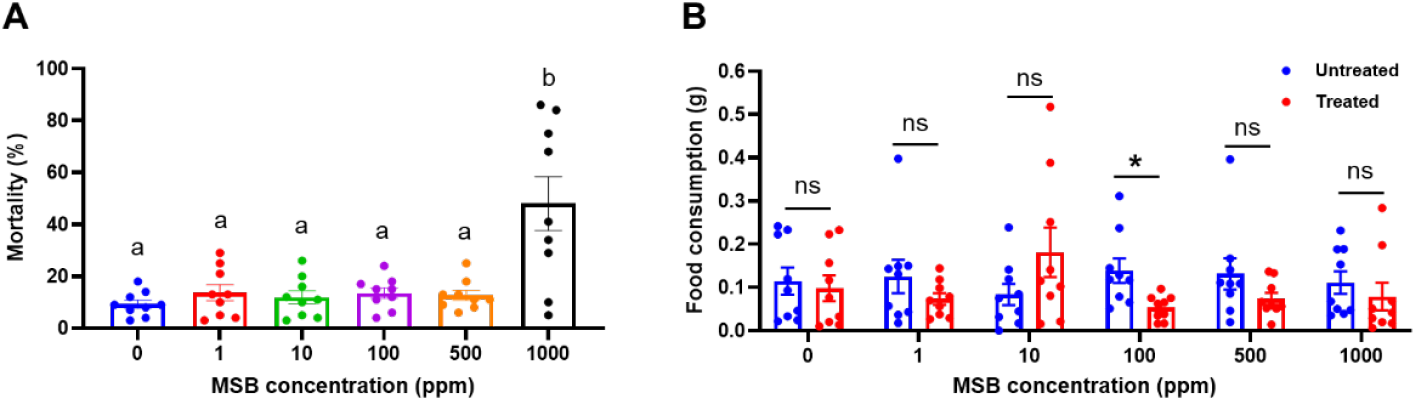
Effects of MSB on termite mortality and feeding in the choice assay. (**A**) Termite mortality 28 days post-treatment. Groups sharing the same letter are not significantly different (GLMM, *df* = 46, *p* > 0.05). (**B**) Wood consumption over 28 days post-treatment (Wilcoxon signed-rank test; ns: not significant (*p* > 0.05); * *p* < 0.05). Bars represent mean ± SE, and dots show individual values.

**Figure 7.**
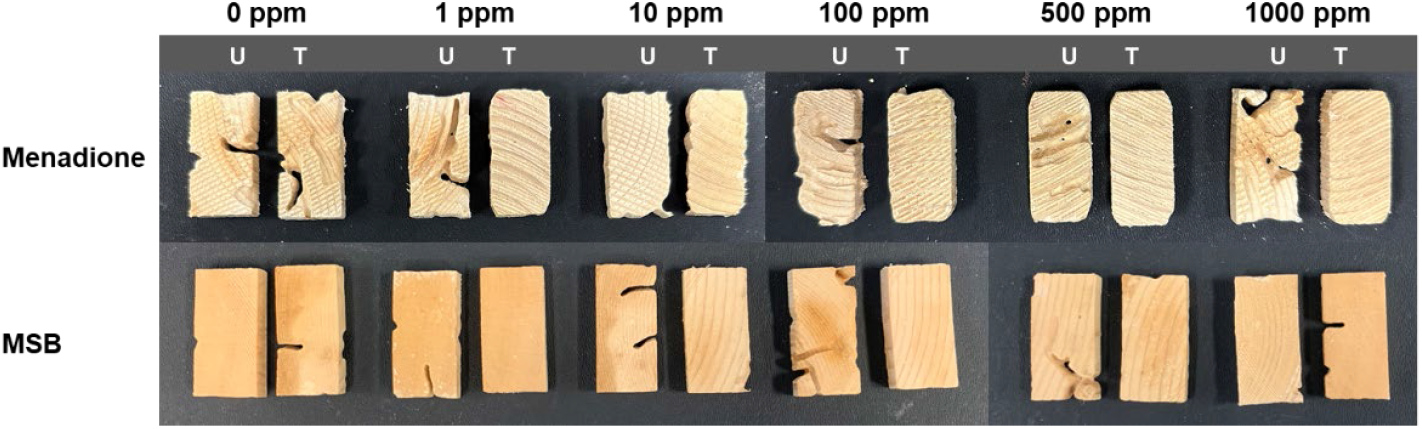
Representative images of wood damage by termites post-menadione and MSB treatments in choice assays. U: untreated; T: treated.

### 4. Discussion

In this study, both menadione and MSB showed toxicity to *C. formosanus* termites, however, the effects varied between the two compounds. While menadione at 1000 ppm caused less than 30% mortality, it reduced feeding by more than 70% compared to the untreated control (Figure 2). In contrast, MSB at the same concentration resulted in much higher mortality (about 75%) but only reduced feeding by 52% (Figure 3). Additionally, the choice assay demonstrated a strong feeding deterrent effect of menadione at 500 and 1000 ppm without causing significant mortality (Figure 5), whereas MSB did not consistently show deterrent activity at the tested concentrations (Figure 6). The results suggest that menadione is more effective than MSB in wood protection due to its feeding deterrence. This compound may be considered for protection against termites from damaging structural wood, wooden furniture, and other cellulose-containing materials such as artwork, pending further testing and optimization.

Compared to our previous study, where soil-treated menadione at 6 ppm triggered significant termite mortality in a no-choice assay and avoidance in a choice assay [31], in this study, a higher concentration of 500 ppm was needed to observe similar effects with wood treatment. This difference is likely due to varying levels of termite exposure to the treatments between the two studies. When only the wood was treated, termites were exposed to the tested chemicals solely through feeding, without contact during tunneling in the substrate sand. Previous research on non-repellent termiticides, such as fipronil, showed that termites can transfer these chemicals to other colony members, leading to increased mortality and potential colony suppression [34-36]. However, such horizontal transfer is less likely to occur with repellent chemicals, and lower mortality is generally expected when termites have access to untreated areas [11], as observed with menadione treatment in this case. Additionally, the diffusion of menadione and MSB is yet to be examined, and it is possible that the chemicals did not penetrate the wood after soaking for 30 seconds. Longer soaking times and application of a second treatment are viable approaches to improve absorption [10]. Although pine wood used in this study is among the most commonly used types of structural wood, treatment optimization requires testing of additional wood species. Also, wood exhibits natural non-uniformity even within the same species, and heterogeneities such as variations in grain patterns, density, and proportions of early- and latewood may influence treatment outcomes.

Menadione may act solely as a feeding deterrent, a repellent, or both to *C. formosanus*, which warrants further investigation through tracking termite behavior. At 500 ppm, menadione treatment significantly reduced but did not completely eliminate food consumption in the choice assay (Figure 5B). Along with the low volatility of menadione, this suggests that direct contact with the chemical was necessary for *C. formosanus* termites to stop feeding, potentially due to perception via their olfactory and gustatory systems [37, 38]. From a practical perspective, however, the outcome is the same: termites can be deterred from consuming the treated wood.

Wood treated with MSB at 1000 ppm exhibited a lethal effect on *C. formosanus* termites in both no-choice and choice assays, but this mortality did not result in significant reductions in wood damage (Figures 3 and 6). MSB may be slow-acting, with termite mortality occurring toward the end of the 28-day assays. The results imply that MSB is non-repellent, or it may require higher doses to exhibit repellency or feeding deterrence, which warrants further investigation. MSB has been previously demonstrated to have insecticidal activity in agricultural pests [27, 39], but its effect on insect feeding has not been reported. While this study did not provide strong evidence supporting the use of MSB as a wood preservative, MSB may be considered for testing as a soil termiticide. Additional studies will be necessary to determine whether this compound is toxic, slow-acting, and non-repellent when applied to soil.

Furthermore, compared to MSB that easily dissolves in aqueous solutions, menadione is less likely to leach out of wood because of its water-insoluble nature. Leachability is an issue associated with water-soluble chemicals, which prevents their effective treatment of wood for exterior use or in contact with ground [40]. For example, borates are widely used wood preservatives due to their broad-spectrum effectiveness and environmentally friendly properties, however, their use is limited to above-ground environments that are protected from weather exposure [40]. Additional coating or fixation of these compounds would be necessary to reduce leaching [40, 41]. Importantly, while water-insolubility is a favorable characteristic of menadione for exterior use, further investigations are needed to optimize its formulation for application and to evaluate its residual effects under varying environmental conditions, such as temperature and light.

Another important future direction is to examine the effects of menadione and MSB on other termites and wood-destroying insects. It is worth noting that insecticide susceptibility could substantially vary between termite species [8]. For example, compared to *Reticulitermes, C. formosanus* is more tolerant to bifenthrin but susceptible to imidacloprid with multi-fold differences in topical LD_50_ [42, 43]. The topical application of chlorfenapyr, which has a similar mode of action to menadione, showed *C. formosanus* to be more susceptible than *R. hesperus* [43, 44]. Additionally, as subterranean termites increase moisture in wood when feeding or foraging, it is also important to evaluate the antifungal properties of potential wood preservatives to ensure their protection against wood-decaying fungi. Since menadione interferes with mitochondrial oxidative phosphorylation [29, 30], a process highly conserved across eukaryotes [45], it will be valuable to further determine whether this compound and its derivatives exhibit broad-spectrum effects and how their efficacy varies among structural pests.

## Supplemental Materials

Figure S1: Original data.

## Author Contributions

Conceptualization, Q.S and R.A.L.; methodology, Q.S. and A.K.; software, A.K.; validation, A.K. and Q.S.; formal analysis, A.K.; investigation, A.K. and M.G.; resources, Q.S.; data curation, A.K.; writing—original draft preparation, A.K.; writing—review and editing, Q.S. and R.A.L; visualization, A.K. and Q.S.; supervision, Q.S.; project administration, Q.S.; funding acquisition, M.G. and Q.S. All authors have read and agreed to the published version of the manuscript.

## Data Availability Statement

The data presented in this study are available in the supplemental materials.

## Acknowledgments

We thank Brendon Davis and Payton Floyed for their assistance with the experiments and termite collection, as well as Dr. Honglin Feng for his comments and discussion about the study.

